# *SUPPRESSOR OF MAX2 1-LIKE 5* promotes secondary phloem formation during radial stem growth

**DOI:** 10.1101/600312

**Authors:** Eva-Sophie Wallner, Virginie Jouannet, Thomas Greb

## Abstract

As a prerequisite for constant growth, plants can produce vascular tissues at different sites in their postembryonic body. In particular, the formation of vascular tissues during longitudinal and radial expansion of growth axes differs fundamentally with respect to its anatomical configuration. This raises the question to which level regulatory mechanisms of vascular tissue formation are shared throughout plant development. Here, we show that, similar as primary phloem formation during longitudinal growth, the cambium-based formation of secondary phloem depends on the function of *SMXL* genes. Using promoter reporter lines, we observe that *SMXL4* and *SMXL5* activities are associated with different stages of secondary phloem formation in Arabidopsis stems and the specific loss of *SMXL5* function results in the absence of secondary phloem. Interestingly, the additional disruption of *SMXL4* activity increases cell proliferation rates in the cambium region without that secondary phloem is formed. Based on genome-wide transcriptional profiling and expression analyses of phloem-related markers we conclude that early steps of phloem formation are impaired in *smxl4;smxl5* double mutants and that additional cambium-derived cells fail in establishing any phloem-related feature. Our results show that molecular mechanisms determining primary and secondary phloem share important features but differ slightly with *SMXL5* playing a more dominant role in the formation of secondary phloem.

## INTRODUCTION

Bodies of higher vascular plants, such as trees, acquire an impressive amount of biomass that solely relies on post-embryonic growth. Continuous formation of organs and tissues is driven by distinct stem cell niches called meristems (Greb and Lohmann 2016). Longitudinal growth of shoots and roots depends on shoot and root apical meristems (SAM, RAM), respectively. These proliferating tissues establish primary shoot and root axes which harbour vascular bundles consisting of two types of primary vascular tissues: The water and nutrient-conducting xylem and the photoassimilate-allocating phloem (Ruonala *et al.* 2017). Naturally, both xylem and phloem are life-essential conduits for long-distance transport that determine plant fitness (De Rybel *et al.* 2016). After establishment of primary growth axes, axes of dicotyledonous species and conifers grow radially with the help of a lateral meristem, called the vascular cambium. The cambium produces secondary phloem and xylem in a bidirectional manner (Campbell and Turner 2017). Therefore, although functionally equivalent, primary and secondary vascular tissues originate from different cellular environments. This raises the question to which extend molecular circuits instrumental for forming primary and secondary vascular tissues are comparable (Jouannet *et al.* 2014).

Formation of primary vascular tissues is extensively studied in root tips of *Arabidopsis thaliana* (Arabidopsis) (De Rybel, et al. 2016) and a large body of literature emphasizes the tight intercellular crosstalk during vascular patterning (Baima *et al.* 2001, Bishopp *et al.* 2011, Carlsbecker *et al.* 2010, Miyashima *et al.* 2019, Smetana *et al.* 2019). Especially, a mutual negative interaction between auxin and cytokinin signalling is essential for establishing domains of xylem and phloem production (Bishopp, et al. 2011, De Rybel *et al.* 2014, Mähönen *et al.* 2006, Ohashi-Ito *et al.* 2014). For the promotion of procambial divisions between both domains, phloem-derived or locally expressed DNA-BINDING ONE ZINC FINGER (DOF) transcription factors promote cell proliferation (Miyashima, et al. 2019, Smet *et al.* 2019). DOFs are counteracted by CLASS III HOMEODOMAIN-LEUCINE ZIPPER (HD-ZIP III) transcription factors which are expressed in early xylem cells where they promote xylem differentiation and instruct cambium organization in a non-cell autonomous manner (Miyashima, et al. 2019, Smetana, et al. 2019).

Except promoting *HD-ZIP III* gene activity, the phloem-borne DOF transcription factor PHLOEM EARLY DOF2 (PEAR) directly activates the *SUPPRESSOR OF MAX2 1-LIKE 3* (*SMXL3*) gene which not only stimulates procambial cell divisions (Miyashima, et al. 2019), but also determines phloem specification (Wallner *et al.* 2017). In this context, *SMXL3* acts together with its closest homologues *SMXL4* and *SMXL5* as key regulators of primary phloem formation (Wallner, et al. 2017). This role is reflected in the absence of primary phloem in *smxl3;4;5* triple mutants, while double mutants show specific defects in (proto)phloem initiation and differentiation within the RAM (Wallner, et al. 2017). Although *SMXL3/4/5* function has not yet been related to other phloem regulators (Anne and Hardtke 2018) and their molecular role is still obscure, early expression of all three genes in phloem precursor cells and their profound role in phloem formation define *SMXL3/4/5* genes as central promoters of phloem formation (Wallner, et al. 2017). These features define *SMXL3/4/5* genes as ideal subjects to compare the mechanisms underlying the development of primary and secondary phloem in different anatomical environments.

Here we report a specific role of *SMXL5* in promoting secondary phloem formation during radial stem growth in *Arabidopsis*. By investigating genome-wide and promoter reporter-based gene expression studies in grafted plants, we observe that secondary phloem formation is disrupted in *smxl5* single and *smxl4;smxl5* double mutants. In contrast to *SMXL4* and *SMXL5*, which are expressed at different stages during secondary phloem formation, *SMXL3* is not expressed during secondary phloem formation in the *Arabidopsis* stem. Based on our observations, we conclude that there are functional similarities and differences within the *SMXL3/4/5* sub-clade and that the formation of primary and secondary phloem shares essential features.

## RESULTS

### *SMXL4* and *SMXL5* promoters are active during secondary phloem formation

*SMXL5* promoter activity marks a cambium sub-domain which is located distally to bifacial cambium stem cells and has been associated with phloem formation during radial plant growth (Shi *et al.* 2019). To see whether there is an association of the primary phloem-promoting *SMXL3/4/5* genes with secondary phloem, we investigated *SMXL3* and *SMXL4* promoter activities in cross sections of stems displaying secondary tissue conformation and compared those activities to the promoter activity of *SMXL5*. Secondary tissue conformation is established by the formation of the interfascicular cambium (IC) between vascular bundles of stems (Altamura *et al.* 2001, Mazur *et al.* 2014, Sanchez *et al.* 2012, Sehr *et al.* 2010). The resulting cambium cylinder produces xylem proximally (towards the stem centre) and phloem distally (towards the stem periphery, Figure 1a). Therefore, vascular tissues found in interfascicular regions are clearly cambium-derived and of secondary nature.

**Figure 1.**
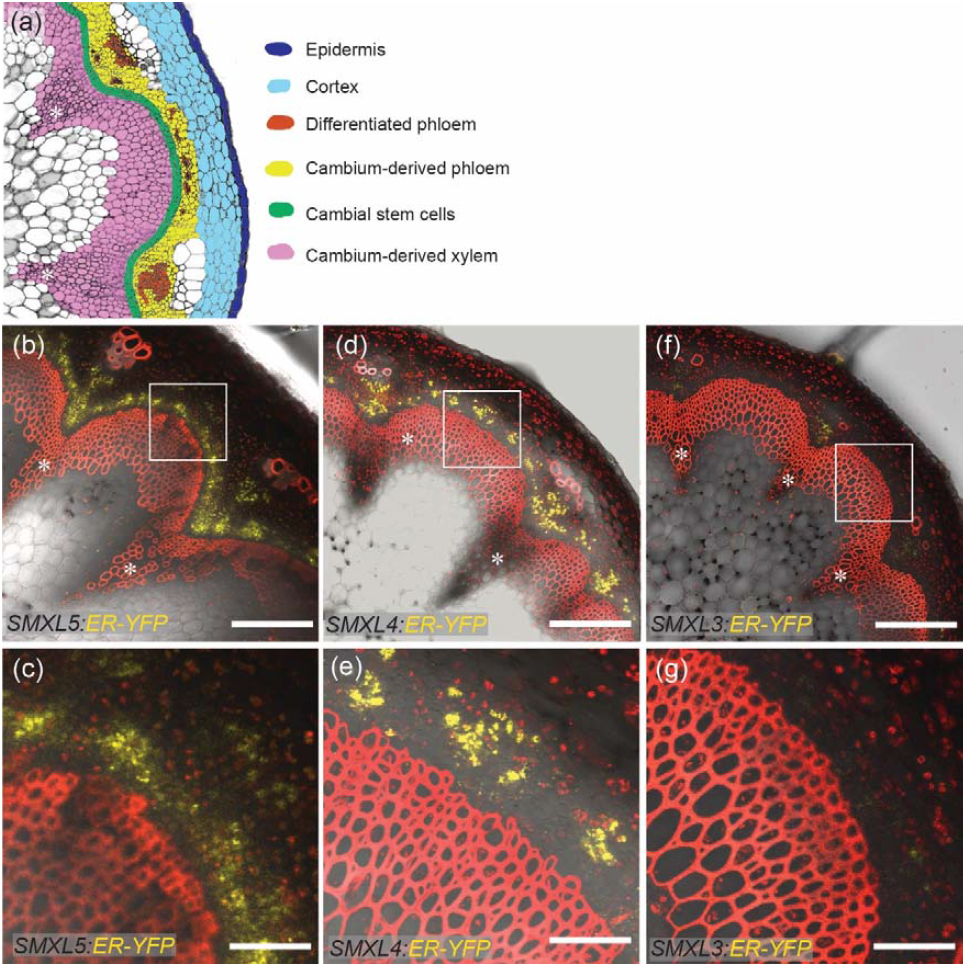
*SMXL4* and *SMXL5* promoters are active in cambium and phloem during radial growth. (a) Schematic representation of a cross section at the stem base. Important tissues are highlighted by different colours as indicated. (b – g) Hand sections of stem bases expressing *SMXL5:ER-YFP* in cambium-derived phloem and differentiated phloem (b, c), *SMXL4:ER-YFP* in differentiated primary and secondary phloem (d,e), and *SMXL3:ER-YFP* in the phloem poles of vascular bundles (f, g). White rectangles mark interfascicular regions shown as close-ups in (c, e, g). YFP signals are depicted in yellow. Stem sections were counterstained with propidium iodide (PI) to visualize xylem in red. Asterisks mark vascular bundles. Scale bars represent 200 µm (b, d, f) or 50 µm (c, e, g).

When analysing transgenic lines expressing endoplasmic reticulum (ER)-localized YFP (YELLOW FLUORESCENT PROTEIN) under the control of the respective *SMXL* promoters, we found *SMXL5:YFP-ER* activity, as expected, in undifferentiated cambium cells and distally from these cells in developing and mature phloem (Figure 1b, c). In comparison, *SMXL4:YFP-ER* activity was observed exclusively in differentiated primary and secondary phloem (Figure 1d, e) whereas only weak *SMXL3:YFP-ER* activity was observed within the primary phloem of vascular bundles (Figure 1f, g). This was in accordance with previous findings of *SMXL3* activity being strong in the seedling root but weak in above-ground tissues (Wallner, et al. 2017). Together, these observations indicated different roles of *SMXL3/4/5* genes in vascular development.

### *SMXL4* and *SMXL5* suppress cambium activity in the stem

Since *SMXL3* activity was not found in secondary phloem of stems, we focused on *SMXL4* and *SMXL5* to determine their roles in secondary phloem formation. To this end, we compared the extension of cambium-derived tissue (CDT) in interfascicular regions of wild type, *smxl4, smxl5* and *smxl4;smxl5* mutants. *smxl4* and *smxl5* single mutants did not differ in CDT extension (Figure 2a-c, e) from wild type plants. However, analysis of the *smxl4;smxl5* double mutant revealed a significant increase in CDT production compared to wild type and the single mutants (Figure 2d, e). Because *smxl4;smxl5* plants showed a weaker growth habit overall (Figure 2f), possibly due to defects in root growth (Wallner, et al. 2017), we investigated stem anatomy after grafting wild type roots to *smxl4;smxl5* shoots and vice versa. Self-grafts of wild type and *smxl4;smxl5* served as controls. Interestingly, *smxl4;smxl5* stems showed an increase in CDT production independently from the root genotype (Figure 3a, b, d, e, f, h, i). In contrast, stem diameter and overall growth of *smxl4;smxl5* shoots supported by a wild type root were fully restored while wild type stems supported by a *smxl4;smxl5* root showed small stem diameter and weak overall growth (Figure 3a-d, j, k). We therefore concluded that deficiency of *SMXL4* and *SMXL5* leads to increased cambium proliferation in stems independently from root or shoot growth.

**Figure 2.**
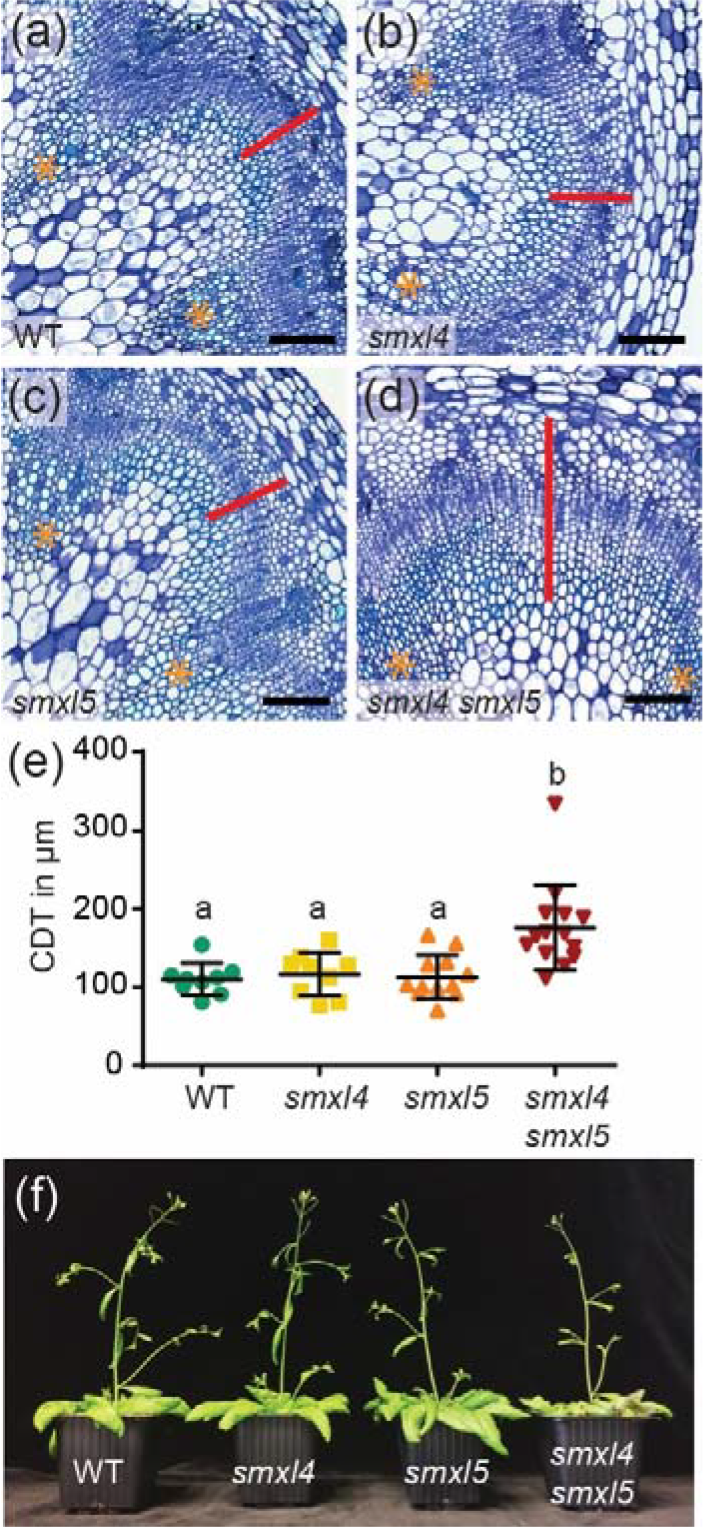
Cambium-derived tissue (CDT) production is increased in *smxl4;smxl5*. (a – d) Representative interfascicular regions of toluidine blue-stained stem base cross sections of wild type (a), *smxl4* (b), *smxl5* (c), and *smxl4;smxl5* (d) are depicted. Yellow asterisks mark vascular bundles. The red line indicates the measured area of cambium-derived tissue (CDT) and follows cell files that originate from cambial stem cells. Scale bars represent 100 µm. (e) Quantification of CDT per genotype. Scatter plots with mean are depicted. Error bars: ± standard deviation (n = 9-13). One-way ANOVA with post-hoc Tukey-HSD (CI 95%) was performed. Statistical differences between sample groups are indicated by letters. (f) Growth habitus of analysed genotypes at a stem height of 15-20 cm.

**Figure 3.**
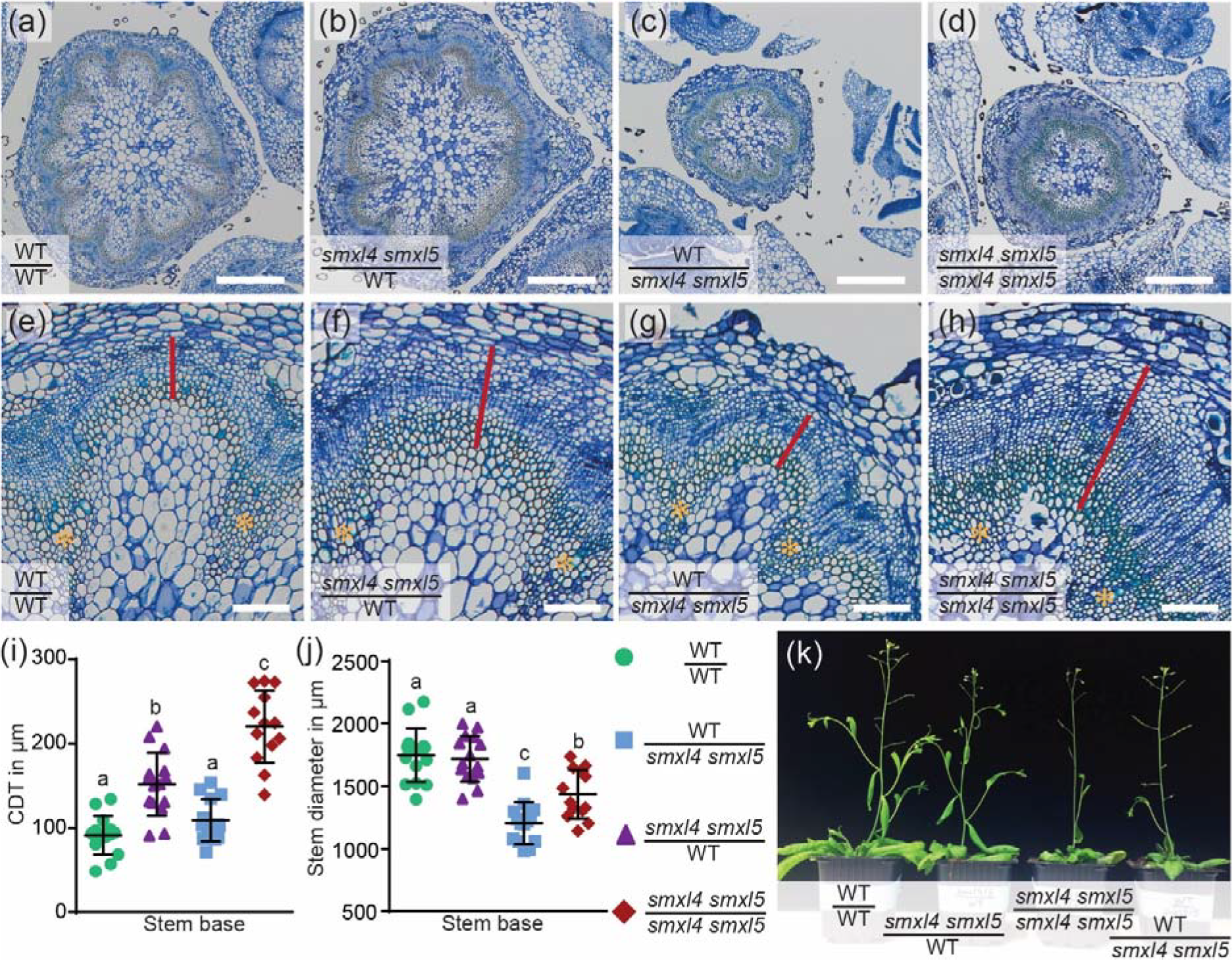
Increase of CDT production in *smxl4;smxl5* is largely independent of the root genotype. (a – d) Toluidine-blue stained cross sections taken at the stem base of 15-20 cm tall grafted *Arabidopsis* plants. The genotypes of shoots (written above the line) and roots (written below the line) of all grafted combinations are indicated in each picture. Scale bars represent 500 µm. (e – h) Close-ups of interfascicular regions of samples described in (a – d). The extensions of CDT are indicated by a red line that follows cambium-derived cell files. Yellow asterisks mark vascular bundles. Scale bars represent 100 µm. (i, j) CDT production (i) and stem diameters (j) of grafted plants. Scatter plots with means are depicted. Error bars: ± standard deviation (n = 13-15). One-way ANOVA with post-hoc Tukey-HSD (CI 95%) was performed in both cases. Wild type shoots grafted on wild type roots are depicted by green cycles, wild type shoots grafted on *smxl4;smxl5* roots are depicted by blue squares, *smxl4 smxl5* shoots grafted on wild type roots are depicted by violet triangles and *smxl4;smxl5* shoots grafted on *smxl4 smxl5* roots are depicted by red rhombi. (k) Growth habitus of grafted plants at a stem height of 15-20 cm.

### *SMXL5* promotes differentiation of secondary phloem

To see whether increased CTD production in *smxl4;smxl5* mutants is associated with altered formation of secondary phloem, we visualized callose, a marker for differentiated sieve elements (SEs) (Barratt *et al.* 2011), in wild type, *smxl4, smxl5* and *smxl4;smxl5* stems as well as in our grafts by aniline staining. These analyses revealed callose deposition in 98.6 % and 94.2 % of the analysed interfascicular regions in wild type and *smxl4* plants, respectively (Figure 4a, b, e, f, i). In contrast, 85.2 % of interfascicular regions analysed in *smxl5* stems were deprived of callose deposition (Figure 4c, g and i). Similarly, 76.1 % of all interfascicular regions in *smxl4;smxl5* stems showed no signs of callose deposition (Figure 4d, h, i). Of note, we obtained similar results when analysing grafted samples. Independently from the root genotype, *smxl4;smxl5* stems were largely deprived of secondary phloem formation within interfascicular regions, while wild type stems supported by a *smxl4;smxl5* root remained unaffected (Figure S1). This indicated that increased CDT production is not accompanied by increased phloem differentiation and that the formation of secondary phloem is indeed *SMXL5*-dependent. Importantly, primary (proto)phloem formation in primary vascular bundles (Figure 4b, c, f, g) and the RAM of *smxl5* or *smxl4* single mutants (Figure S2) resembled wild type plants. Thus, we concluded that molecular mechanisms determining primary and secondary phloem shared important features but differed slightly with *SMXL5* playing a more dominant role in the formation of secondary phloem. Moreover, disruption of *SMXL4* function in a *SMXL5*-deficient background leads to increased proliferation of the cambium in stems.

**Figure 4.**
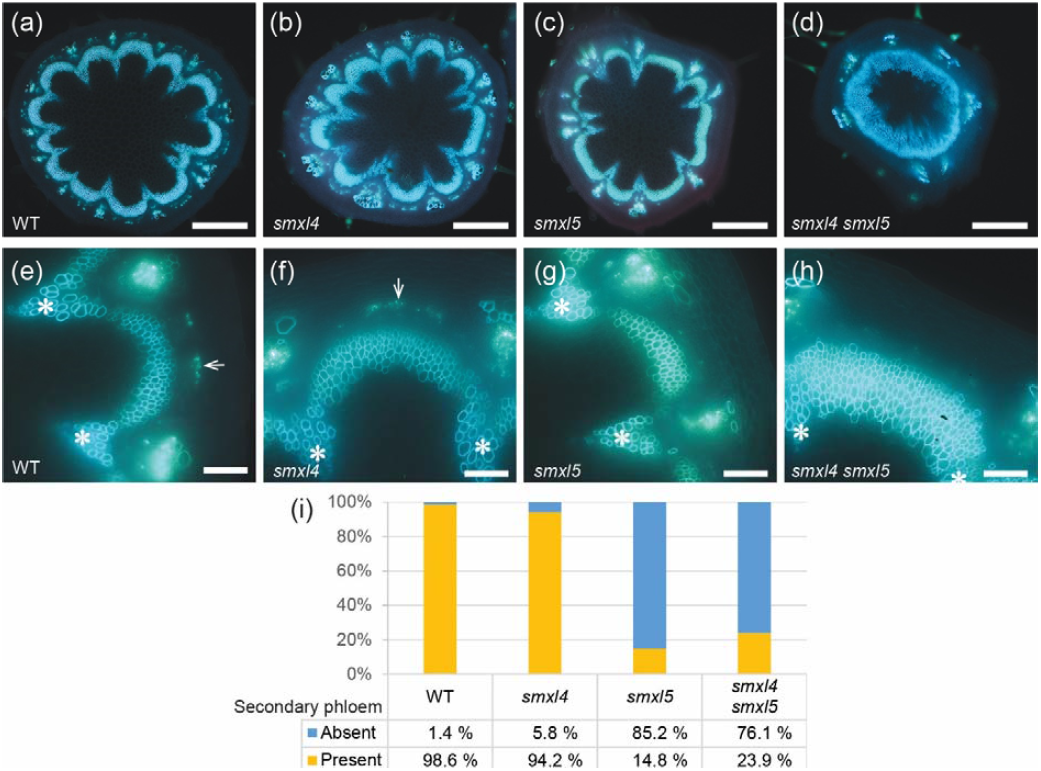
Secondary phloem is reduced in *smxl5* and *smxl4;smxl5* (a – d) Aniline blue stained stem base cross sections of wild type (a), *smxl4* (b), *smxl5* (c) and *smxl4;smxl5* (d). Aniline-stained callose/phloem in blue/green and in light blue. Scale bars represent 500 µm. (e – h) Close-ups of interfascicular regions are depicted for samples described in (a – d). Callose deposition in secondary phloem is marked by arrows in E and F and is absent in G and H. White asterisks mark vascular bundles. Scale bars represent 100 µm. (i) Average percentages of secondary phloem absence (blue) and presence (yellow) are depicted in a stacked histogram (n = 19-23).

### Phloem markers are absent in interfascicular regions of *smxl5* stems

To characterize the anatomical alterations in *smxl4;smxl5* mutant stems, we compared the transcriptomes of *smxl4;smxl5* stem bases supported by a wild type root to those of wild type self-grafts. Application of an adjusted p-value cut-off of 0.01 and a fold change > 1.5 and < 0.5, identified 1343 downregulated and 273 upregulated genes in *smxl4;smxl5* stems, respectively (Figure 5a and Table S1). Since we were especially interested in vasculature-related responses within the *smxl4;smxl5* mutant stem, we first compared our datasets to a set of 280 vascular genes described previously (Endo *et al.* 2014). From those 280 genes, we found 98 to be downregulated in *smxl4;smxl5*, while only 13 were upregulated (Figure 5a). As expected, the downregulated fraction contained several important phloem regulators, such as *ALTERED PHLOEM DEVELOPMENT (APL)* (Bonke *et al.* 2003), *SIEVE ELEMENT OCCLUSION RELATED 1 (SEOR1)* (Froelich *et al.* 2011), *BREVIS RADIX (BRX)* (Mouchel *et al.* 2004, Rodriguez-Villalon *et al.* 2014), *BARELY ANY MERISTEM 3 (BAM3)* (Depuydt *et al.* 2013), *NAC DOMAIN CONTAINING PROTEIN 86 (NAC086)* (Furuta *et al.* 2014), *JULGI1* and *JULGI2* (Cho *et al.* 2018) or *CELLULOSE SYNTHASE (CESA)* subunits *4* and *8*. *CESA4* and *CESA8* are important during secondary cell wall formation of differentiating SEs (Taylor *et al.* 2003). To see whether phloem development is disrupted at a distinct state, we next compared our identified genes with four groups of genes (modules) associated with four different stages of phloem formation (Kondo *et al.* 2016). As a result, we found strong enrichment of our downregulated genes in all four modules. In contrast, only one of the upregulated genes (*AT5G04310*) was part of module IV (Figure 5b and Table S1). Collectively, our RNA sequencing data showed downregulation of a large spectrum of phloem-related genes reinforcing our finding of impaired secondary phloem differentiation in *smxl4;smxl5*. Moreover, the observation that, in addition to ‘late’ modules, a large fraction of the genes comprising module I was downregulated in *smxl4;smxl5* mutants implying that already early events of phloem formation depend on the function of *SMXL4* and *SMXL5*.

**Figure 5.**
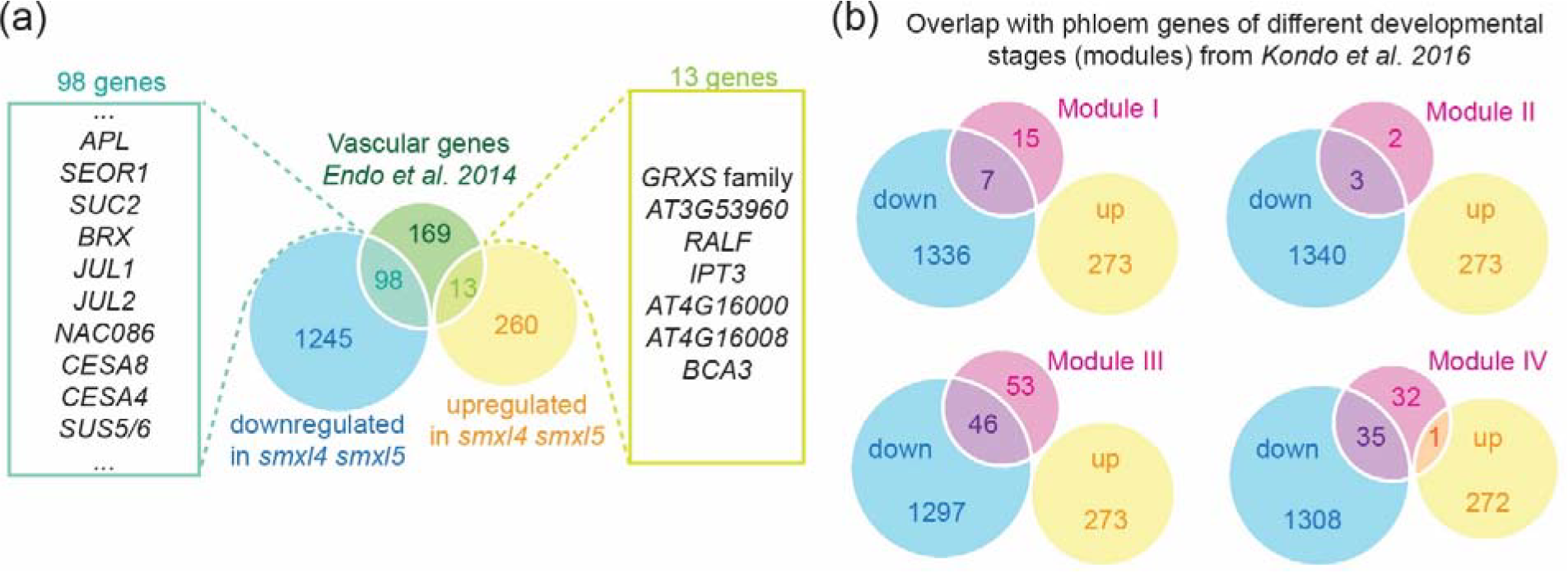
Phloem-related genes are downregulated in secondary *smxl4;smxl5* stems grafted onto wild type roots (a) Venn diagram comparing up- and downregulated genes in *smxl4;smxl5* (with an adjusted p-value < 0.01 and a fold change > 1.5 for downregulated genes and < 0.5 for upregulated genes, respectively) to the dataset of vascular genes (Endo et al. 2014). (b) Venn diagrams comparing the same up- and downregulated genes in *smxl4;smxl5* to genes expressed during different stages of phloem formation (Kondo et al. 2016). Module I contains early, and module IV late genes upregulated upon phloem induction.

Interestingly, the upregulated fraction mostly included genes involved in stress responses, such as *GLUTAREDOXIN S* family members (GRXS) (Moseler *et al.* 2015) or *RAPID ALKALIZATION FACTOR (RALF)* (Sharma *et al.* 2016). Despite more CDT was produced in *smxl4;smxl5* (Figure 2, Figure 3), though, genes associated with cambium stem cells, such as *PHLOEM INTERCALATED WITH XYLEM* (*PXY*)*, SMXL5, WUSCHEL HOMEOBOX RELATED 4* (*WOX4*), *AINTEGUMENTA* (*ANT*) or *HIGH CAMBIAL ACTIVITY 2* (*HCA2*) (Etchells *et al.* 2013, Guo *et al.* 2009, Shi, et al. 2019, Smetana, et al. 2019), were not significantly changed (Table S1). These observations suggested that the number of cambium stem cells was not increased in *smxl4;smxl5* mutants and that additional cambium-derived cells acquired an identity unrelated to vascular cell fates.

To confirm these assumptions, we crossed cambium and phloem markers into *smxl4, smxl5* and *smxl4;smxl5* plants and investigated activity patterns and intensities by confocal microscopy. Promoter activities of *PXY:mTurquoise-ER* and *SMXL5:VENUS-ER* mark the proximal and distal cambium domains, respectively (Shi, et al. 2019), and we expected an increase especially in the number of *SMXL5*-positive cells if the additional cells were arrested at early stages of phloem development. However, the extension of the cambium domains marked by *PXY:mTurquoise-ER* and *SMXL5:VENUS-ER* did not differ between *smxl4;smxl5*, wild type and the *smxl* single mutants in non-grafted or grafted plants (Figure 6 and Figure S3). This confirmed our expectation that the actual stem cell pool including the phloem-forming distal cambium domain was not altered in *smxl4;smxl5*. Moreover, impaired phloem formation in *SMXL5*-deficient plants did not affect cambium organization.

**Figure 6.**
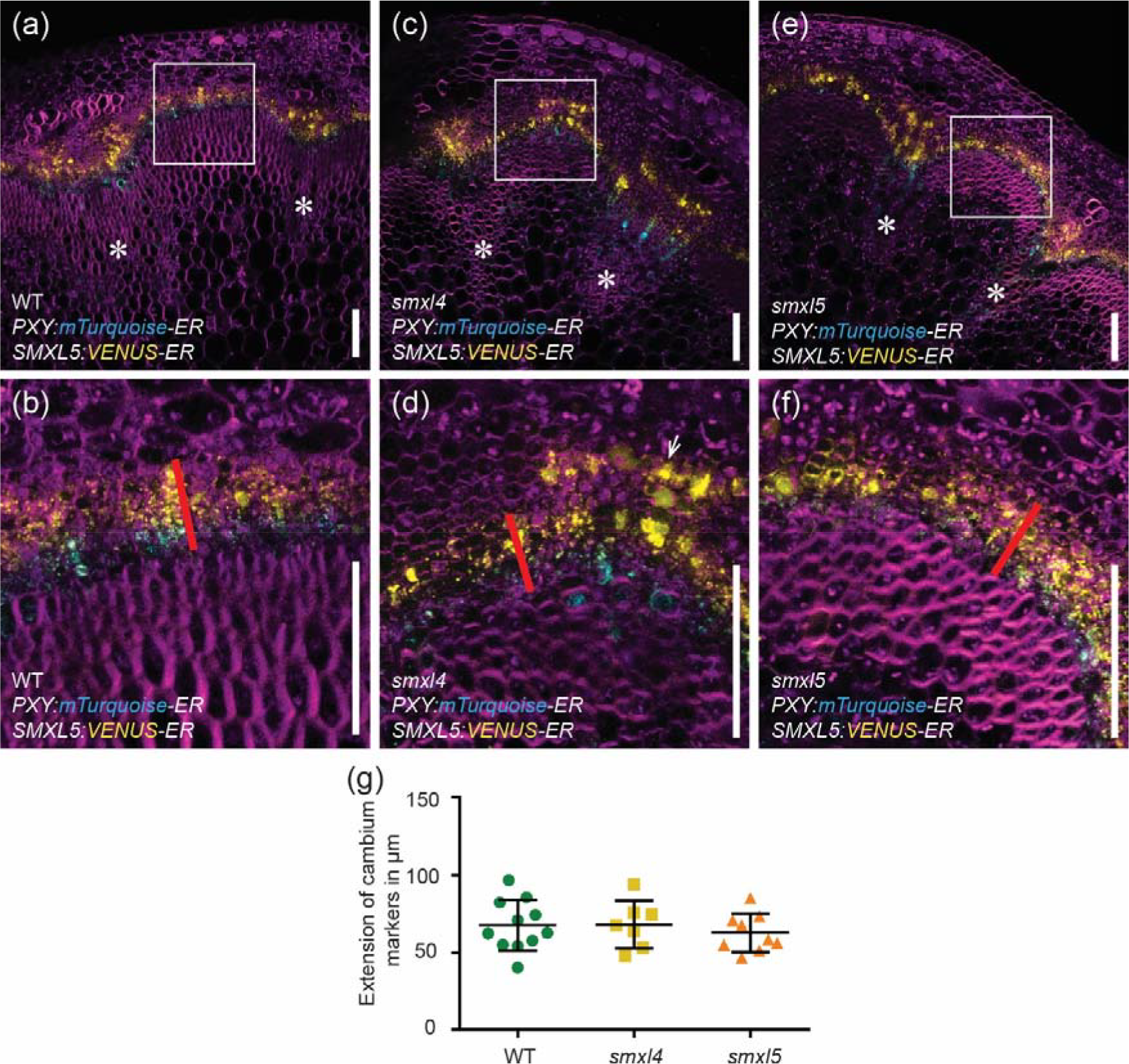
Extension of cambium markers is comparable between genotypes (a – f) Stem base cross sections of *PXY:mTurquoise-ER*;*SMXL5:VENUS-ER* carrying plants were analyzed by confocal microscopy. Depicted are representative stem sections of wild type (a, b), *smxl4* (c, d), and *smxl5* (e, f). White rectangles in (a, c, e) mark interfascicular regions shown as close-ups in (b, d, f). Cell walls were counterstained with DirectRed23 (in magenta). mTurquoise and VENUS signals are shown in cyan and yellow, respectively. White arrows point to marker activity in secondary phloem. White asterisks mark interfascicular bundles. Red lines show measured extension of cambium marker-derived fluorescent signals. Scale bars represent 100 µm. (g) Extension of cambium marker-derived fluorescent signals was compared between genotypes. Scatter plots with means are depicted. Error bars: ± standard deviation (n = 7-11 individual plants per genotype). One-way ANOVA with post-hoc Bonferroni (CI 95%) was performed. No significant differences were detected between sample groups.

To determine the state of cells present distally from cambium stem cells, we analysed promoter-reporter activities of *SMXL4* and *SIEVE ELEMENT OCCLUSION 2* from *Medicago truncatula* (*MtSEO2*) as early and late phloem markers, respectively (Froelich, et al. 2011, Knoblauch and Peters 2010, Wallner, et al. 2017). Interestingly, neither marker was active in IC regions of secondary *smxl5* and *smxl4;smxl5* stems grafted onto wild type roots (Figure 7 and Figure S4). In contrast, the activities of *SMXL4* and *MtSEO2* promoter-reporters were abundant in secondary phloem of wild type and *smxl4* and detected in primary phloem of all analysed genotypes (Figure 7 and Figure S4). This indicated that differentiation of secondary phloem and particularly SEs derived from the cambium in Arabidopsis stems was promoted by *SMXL5*. In addition, we did not find indications that additional cambium-derived cells in *SMXL4/5*-deficient plants established phloem-related signatures.

**Figure 7.**
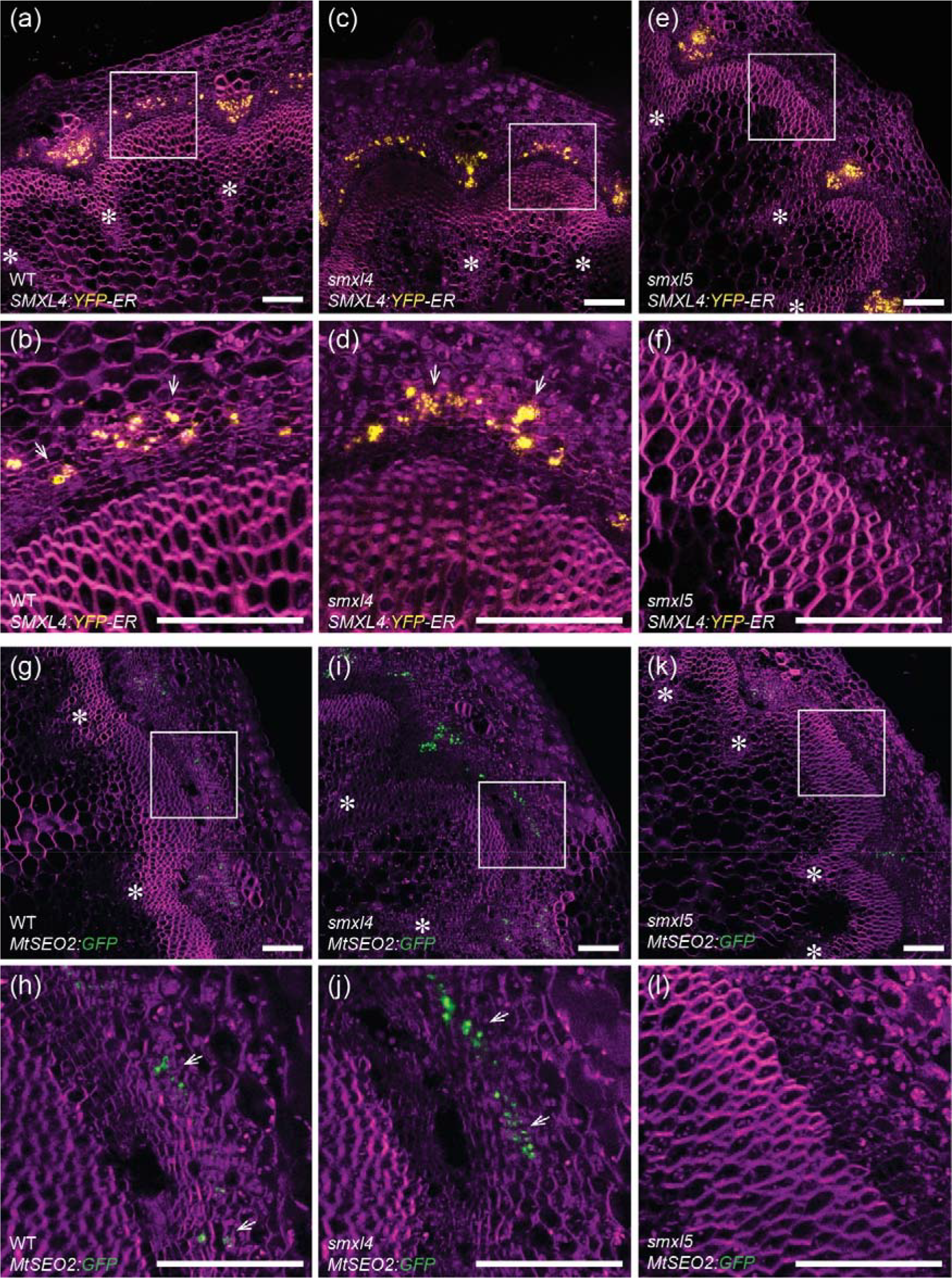
Phloem markers are absent in interfascicular regions of *smxl5* (a – l) Stem base cross sections of marker lines *SMXL4:YFP-ER* (a – f) and *MtSEO2:GFP-ER* (g – l)) were analyzed by confocal microscopy. Depicted are representative stem sections of wild type (a, b, g, h), *smxl4* (c, d, i, j), and *smxl5* (e, f, k, l). White rectangles in (a, c, e, g, I, k) mark interfascicular regions shown as close-ups in (b, d, f, h, j, l). Cell walls were counterstained with DirectRed23 (in magenta). YFP and GFP signals are shown in yellow and green, respectively. White asterisks mark interfascicular bundles. White arrows point to marker activity in secondary phloem. Scale bars represent 100 µm.

## DISCUSSION

Growth of vascular plants, which represent the major source of food and biomass on earth (Nieminen *et al.* 2012), requires a fine-tuned balance between stem cell maintenance, cell divisions and differentiation (Hong *et al.* 2015, Sozzani and Iyer-Pascuzzi 2014). Despite their fundamental importance, molecular mechanisms behind vascular development are still not fully understood. Here we identified a dual role of *SMXL5* in promoting secondary phloem formation and, together with *SMXL4*, in suppressing CDT production.

*SMXL4* and *SMXL5* fulfil roles as major phloem regulators across different organs and growth stages (Wallner, et al. 2017). By using the SE-specific promoter-reporter *MtSEO2:GFP-ER* we determined that SE formation during secondary phloem formation is largely dependent on *SMXL5*. Moreover, more cells were produced by the cambium in *smxl4;smxl5* double mutants, but cells generated at the distal phloem side failed to establish any phloem-related features. This finding is consistent with previous observations: Differentiation, including enucleation, of SEs is absent in the RAM of *smxl4;smxl5* double mutants and primary phloem within vascular bundles of *smxl4;smxl5* stems accumulated undifferentiated, still nucleated cells (Cho, et al. 2018, Wallner, et al. 2017). Interestingly, we observed defects in secondary phloem formation already in *smxl5* single mutants, which showed normal SE differentiation within the RAM and the primary phloem of stems. Given the differences in expression domains and levels of *SMXL3/4/5* gene activities in secondary stems, we propose specialized roles of those otherwise redundantly acting phloem regulators during radial plant growth.

Comparing *SMXL3/4/5* function in phloem formation in the RAM and the IC may shed light on cambium regulation. Already in the embryo the quiescent centre (QC) gives rise to four types of so-called initials (Scheres 2007, Sozzani and Iyer-Pascuzzi 2014). Those are root stem cells with pre-determined fate towards e.g. cortex, vasculature, epidermis or columella (Sozzani and Iyer-Pascuzzi 2014). They are maintained by the otherwise quiescent QC and divide asymmetrically into a stem cell and a daughter cell (De Rybel, et al. 2016, Scheres 2007). Detached from the organizing QC, the daughter cells start to differentiate into specific tissue types (Sozzani and Iyer-Pascuzzi 2014). Phloem daughter cells undergo important anticlinal and tangential cell divisions to initiate more specific cell lineages in a highly controlled spatio-temporal manner (Bonke, et al. 2003, Scacchi *et al.* 2010). In *smxl4;smxl5* mutants, we found the second tangential cell division within the phloem cell lineage to be delayed (Wallner, et al. 2017). Consequently, more cells were produced that failed in differentiating into phloem cells (De Rybel, et al. 2016, Wallner, et al. 2017). We made a similar observation for the cambium: More cambium-derived cells were produced and, towards the distal cambium side, those cambium descendants failed to differentiate. As in the RAM, the nature of these cells is still unknown. Neither the proximal cambium marker *PXY:mTurquoise-ER* nor the distal cambium marker *SMXL5:VENUS-ER* showed an extended activity within *smxl5* or *smxl4;smxl5* stems and other phloem-related markers were fully absent. We hypothesize that those cambial daughter cells are stuck in an undifferentiated state, similar to cells in the RAM of *smxl4;smxl5* mutants.

Exploring the roles of *SMXL* genes in secondary phloem formation, we found indications that *SMXL* genes belong to the earliest players in phloem formation. We found genes downregulated in *smxl4;smxl5* which belong to the earliest genes transcriptionally activated during phloem induction (Module I) (Kondo, et al. 2016). Of note, *SMXL5* is also part of genes in the Module I (Kondo, et al. 2016), which highlights the importance of *SMXL5* during early stages of phloem formation. Overall, we discovered that *SMXL5* plays a key role in a so far unknown regulatory network instructing secondary phloem formation. Therefore, our observations support the conclusion that molecular mechanisms acting in primary and secondary phloem share important features.

## MATERIAL AND METHODS

### Plant material and growth conditions

*Arabidopsis thaliana* (L.) Heynh. Col-0 plants were used in all cases. Genetic analysis was performed with mutant alleles *smxl5-1* (*SALK_018522*) and *smxl4-1* (*SALK_037136*). Genotyping was done by PCR using primers as described previously (Wallner, et al. 2017). Before sowing seeds directly on soil, they were vapour sterilized for 2 h in a sealed container by Hydrogen chloride, which was produced by mixing 40 ml 13% sodium hypochlorite with 1.2 ml 30% hydrochloric acid. For breaking seed dormancy and providing contemporaneous germination, seeds were stratified on soil at 4 °C for 3 days in the dark. Plants were first grown in short day (SD) conditions (8 h light and 16 h dark) with 60 % humidity and 22 °C for 3 weeks and then transferred to long day (LD) conditions (16 h light and 8 h dark) at 22 °C and 60 % humidity.

For grafting experiments and root studies, seeds were sterilized by 70 % ethanol supplemented with 0.2 % Tween-20 for 15 min, washed twice with 100 % ethanol and air dried under sterile conditions. Seeds were sown on Murashige and Skoog (MS) medium-plates containing 0.8 % phyto agar and 1 % sucrose. Seeds were stratified on plates for 3 days at 4°C in the dark. Seedlings were grown in SD conditions at 22°C for 5 days prior to grafting. Freshly grafted seedlings were kept for 6 days on wetted filter paper in sterile conditions to ensure reconnection of the vasculature and then transferred for another 7 days on MS-plates for recovery. Well-connected grafts were transferred to soil and kept in SD for another 3 days. After 21 days (3 weeks) plants were transferred to LD conditions.

### Grafting

Grafting was performed on 5 day-old seedlings grown in SD as previously described (Melnyk and Meyerowitz 2015).

### Generation of plasmids and transgenic lines

Plasmids *SMXL3:YFP-ER, SMXL4:YFP-ER* and *SMXL5:YFP-ER* were generated via a two-step cloning procedure using the vector *pGreen0229* (Hellens *et al.* 2000). The exact procedure was described previously (Wallner, et al. 2017). Construction of the double-marker *PXY:mTurquoise-ER;SMXL5:VENUS-ER* was done by using the Greengate system as described in (Shi, et al. 2019). Col-0 seeds transformed with *MtSEO2:GFP-ER* were obtained from Michael Knoblauch (Froelich, et al. 2011).

### Microscopy and tissue staining

For analysing reporter activities or aniline staining in stems, the stem base was embedded in small polystyrol blocks and hand sections of approximately 0.5 mm were taken by a razor blade (Wilkinson Sword). Reporter activities were analysed by a Leica TCS SP5II (Leica) system equipped with the Leica Application Suite X software. Prior to imaging, fresh stem sections were counterstained for 5 min by 5 µg/ml propidium iodide (PI, Merck) dissolved in tap water or 0.01 % Direct Red 23 (Sigma-Aldrich) dissolved in 1× PBS. mPS-PI staining of roots was carried out as described before (Truernit *et al.* 2008, Wallner, et al. 2017). YFP (yellow fluorescent protein) was excited by an argon laser at 514 nm and the emission detected at 520-540 nm. PI was excited at 561 nm (DPSS laser) and detected at 590-690 nm. Aniline staining was performed as described previously (Schenk and Schikora 2015, Wallner, et al. 2017). Stained stem sections were analysed using the epifluorescence microscope Zeiss Axio Imager.M1 (Carl Zeiss). Fluorescence of aniline was excited by the UV lamp and emission was collected using the DAPI filter.

### Histological analyses

1-cm stem fragments just above the uppermost rosette leaf and 1-cm fragments including the hypocotyl and the uppermost root region were harvested from 15 – 20 cm tall plants and infiltrated by 70 % ethanol for at least 3 days. Samples were processed using the Tissue dehydrator LOGOS Microwave Hybrid Tissue Processor (Milestone) and the LEICA ASP200S (Leica). After embedding in paraffin, 10 μm sections were produced using a microtome, deparaffinised, stained with 0.05% toluidine blue (Applichem), and fixed by Entellan (Merck). Slides were scanned using a MIRAX Slide Scanner (3DHistech) or the Axio Imager.M1 microscope (Carl Zeiss). Scans were analysed using the Pannoramic Viewer (Histech) or ImageJ 1.49d (Schindelin *et al.* 2012).

### RNA extraction and sequencing

RNA was extracted from 1-cm stem segments including the stem base. Three stem segments were pooled into one biological replicate. Tree biological replicates were collected per genotype. RNA was extracted by phenol/chlorophorm as described (Mallory and Vaucheret 2010) and treated with RNase-free DNase (Fermentas). Single reads of 50 nucleotides in length were sequenced using HiSeqV4 SR50. Reads were aligned to the *Arabidopsis* genome (TAIR10) using TopHat2 v2.0.14 (Kim *et al.* 2013) and the statistical analysis was done using the DESeq2 package from the R/Bioconductor software (Love *et al.* 2014). Mutant genotypes were compared to wild type by applying a stringency of adjusted p-value < 0.01. Data processing included the use of VENNY (Oliveros 2007) and alignment to The Arabidopsis Information Resource (TAIR)-databases (Huala *et al.* 2001) by VirtualPlant 1.3 (Katari *et al.* 2010).

### Statistical analysis

Statistical analyses were performed using IBM SPSS Statistics for Windows, Version 23.0. Armonk, NY: IBM Corp. Means were calculated from measurements with sample sizes as indicated in the respective figure legends. Error bars represent ± standard deviation. All analysed datasets were prior tested for homogeneity of variances by the Levene statistic. Statistically different groups were determined by a One-way ANOVA with a confidence interval (CI) of 95 %. If variances were homogeneous (equal), a post-hoc Bonferroni test was applied for datasets n < 5 and a post-hoc Tukey HSD test was applied for datasets with n > 4. For comparison of two datasets a Student’s *t*-test (for equal variances) was performed. Graphs were generated in GraphPad Prism version 6.01 (GraphPad Software), or Excel (Microsoft).

### Accessing RNA sequencing data

Raw data discussed in this publication have been deposited in NCBI’s Gene Expression Omnibus (Barrett *et al.* 2013) and are accessible through GEO Series accession number GSE129190 at https://www.ncbi.nlm.nih.gov/geo/.

## Acknowledgements

This work was supported by the Deutsche Forschungsgemeinschaft (DFG) through the grant GR2104/4-1, the SFB873, and a Heisenberg Professorship [GR2104/5-1] to T.G.. Library preparation and next-generation-sequencing was performed at the Deep Sequencing Core Facility provided by the Exzellenzcluster CellNetworks at the University of Heidelberg (http://www.cellnetworks.uni-hd.de/483065/Deep_Sequencing_Core_Facility).

## Author Contributions

Conceived and designed the experiments: EW, TG. Performed the experiments: EW Analysed the data: EW, VJ. Wrote the paper: EW, TG.

## Conflict of Interest

The authors declare that they have no conflict of interest.

---

**Figure S1.**
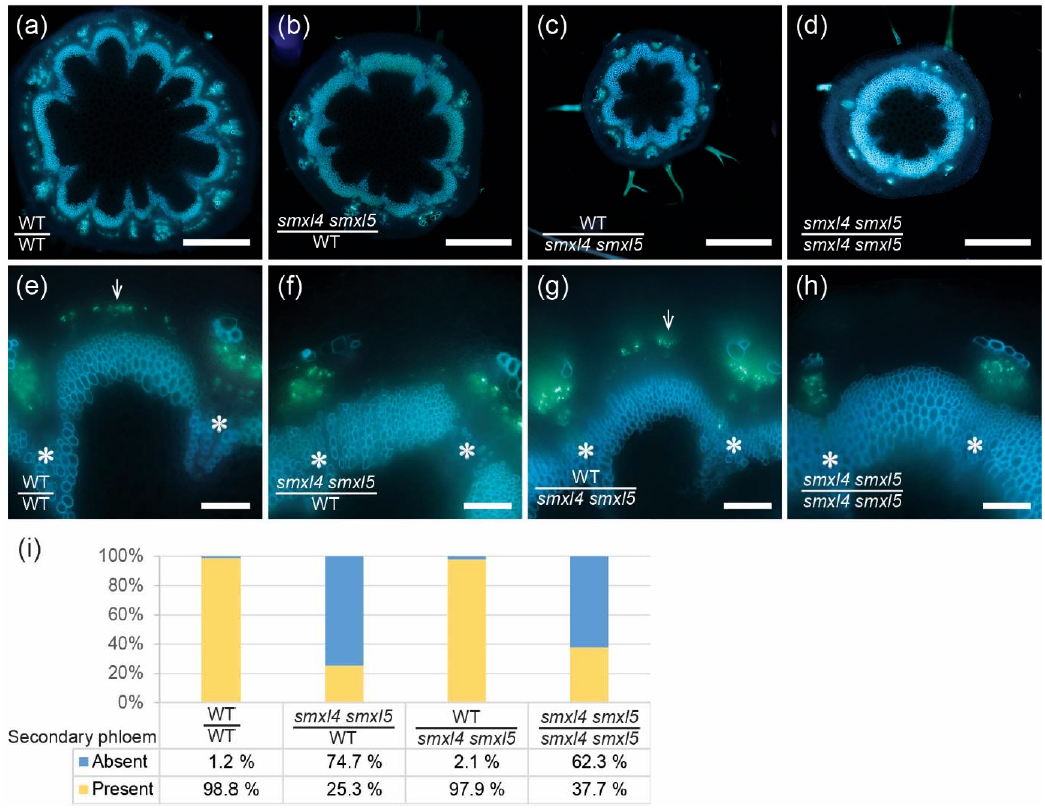
Differentiation of secondary phloem is impaired in *smxl4;smxl5* stems independently of the root genotype. (a – d) Aniline blue stained cross sections taken at the stem base of all grafted combinations are shown as indicated in the pictures. Xylem autofluorescence is light blue and aniline stained callose/phloem green. Scale bars represent 500 µm. (e – h) Close-ups of interfascicular regions of samples described in (a – d). Xylem autofluorescence is light blue and aniline stained callose/phloem green. Callose deposition in secondary phloem is marked by arrows in (e) and (g) and is absent in (f) and (h). White asterisks (*) mark vascular bundles. Scale bars represent 100 µm. (i) Visual representation of the average percentages of interfacicular regions with (e, g) or without (f, h) secondary phloem (n = 10-15). Results could be reproduced in an independent repetition (n = 6-12).

**Figure S2.**
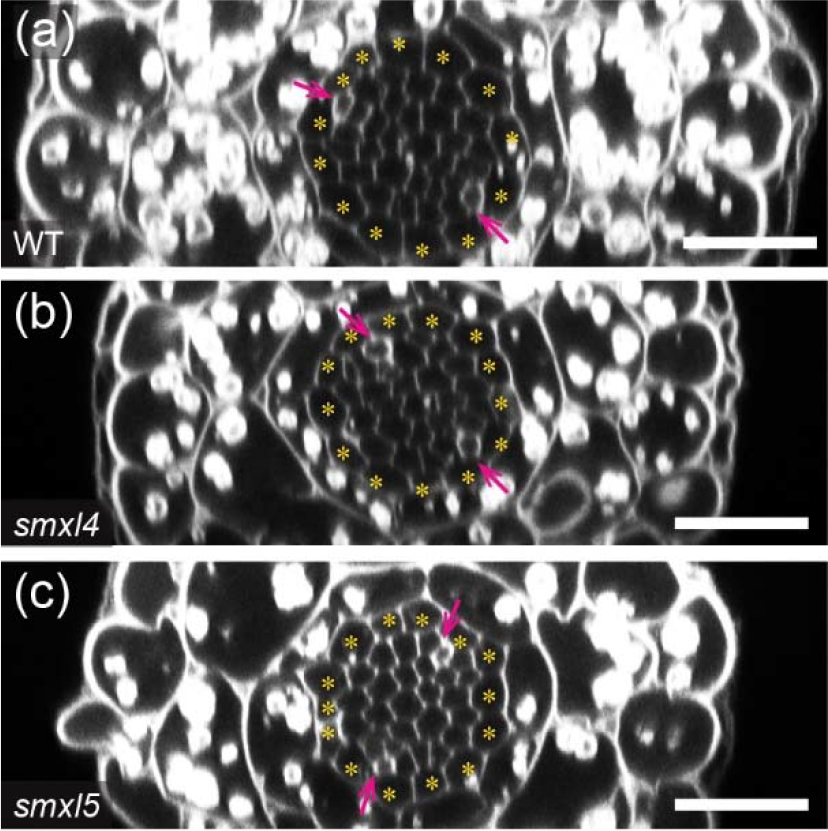
Phloem formation is not affected in *smxl5* roots (a – c) Optical cross sections taken 200 µm proximal to the quiescent centre of two day-old mPS-PI stained root tips from wild type (a), *smxl4* (b) and *smxl5* (c) plants are shown. Yellow asterisks mark pericycle cells. Pink arrows point to differentiating SEs that are indicated by enhanced PI staining. Scale bars represent 20 µm.

**Figure S3.**
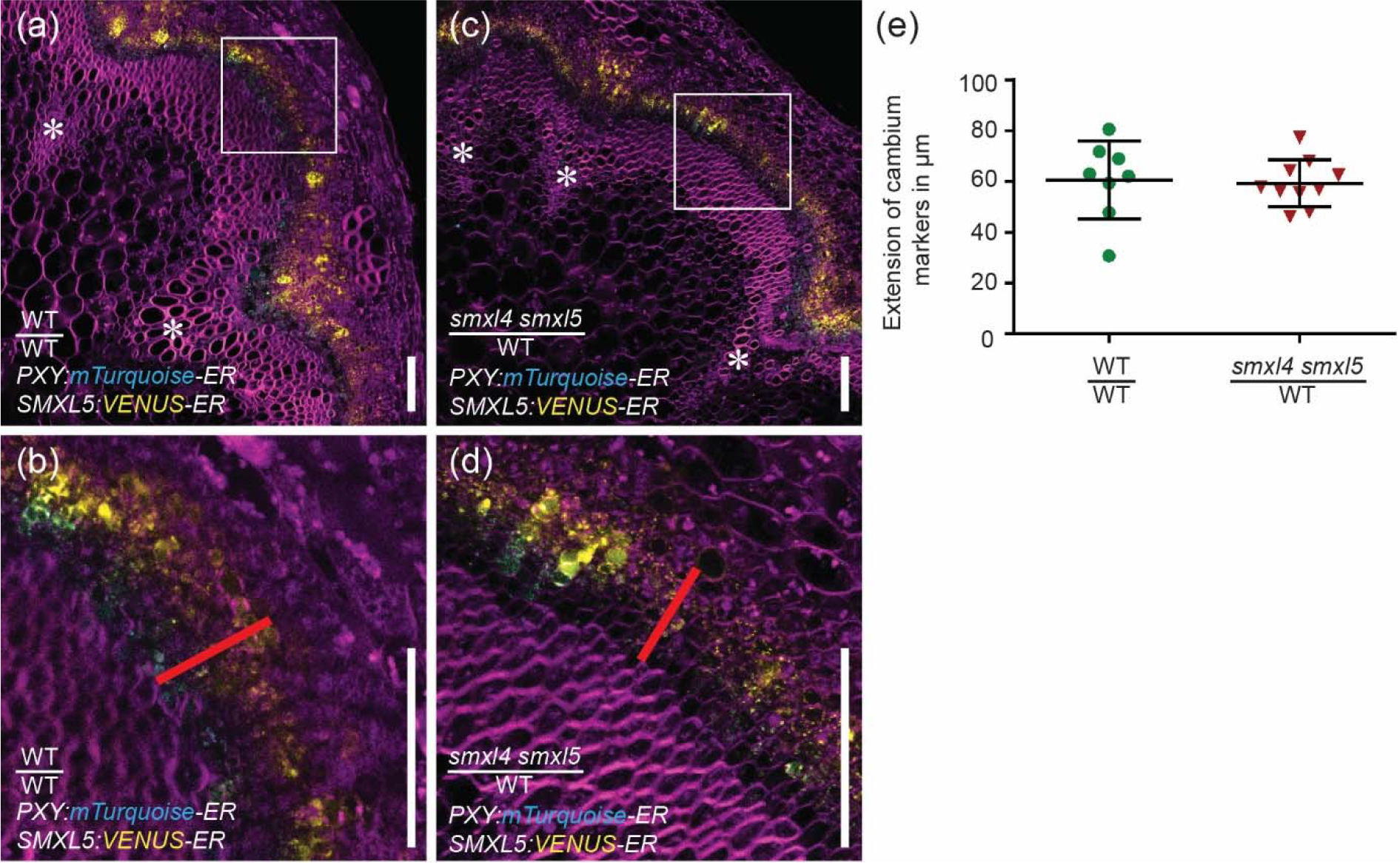
Extension of cambium markers is comparable between wild type and *smxl4;smxl5* (a – d) Stem base cross sections from wild type shoots grafted onto wild type roots (a, b) and from *smxl4;smxl5* shoots grafted onto wild type roots (c, d) expressing *PXY:mTurquoise-ER*;*SMXL5:VENUS-ER* were analyzed by confocal microscopy. Cell walls were counterstained by DirectRed (magenta). White rectangles in (a) and (c) mark interfascicular regions shown as close-ups in (b) and (d). mTurquoise and VENUS signals are shown in cyan and yellow, respectively. White asterisks mark interfascicular bundles. White arrows point to marker activity in secondary phloem. Red lines show measured extension of cambium marker-derived fluorescent signals. Scale bars represent 100 µm. (e) Extension of cambium marker-derived fluorescent signals was compared between genotypes. Scatter plots with means are depicted. Error bars: ± standard deviation (n = 8-10 individual plants per genotype). Student’s *t*-test (CI 95%) was performed. No significant differences were detected between the two sample groups.

**Figure S4.**
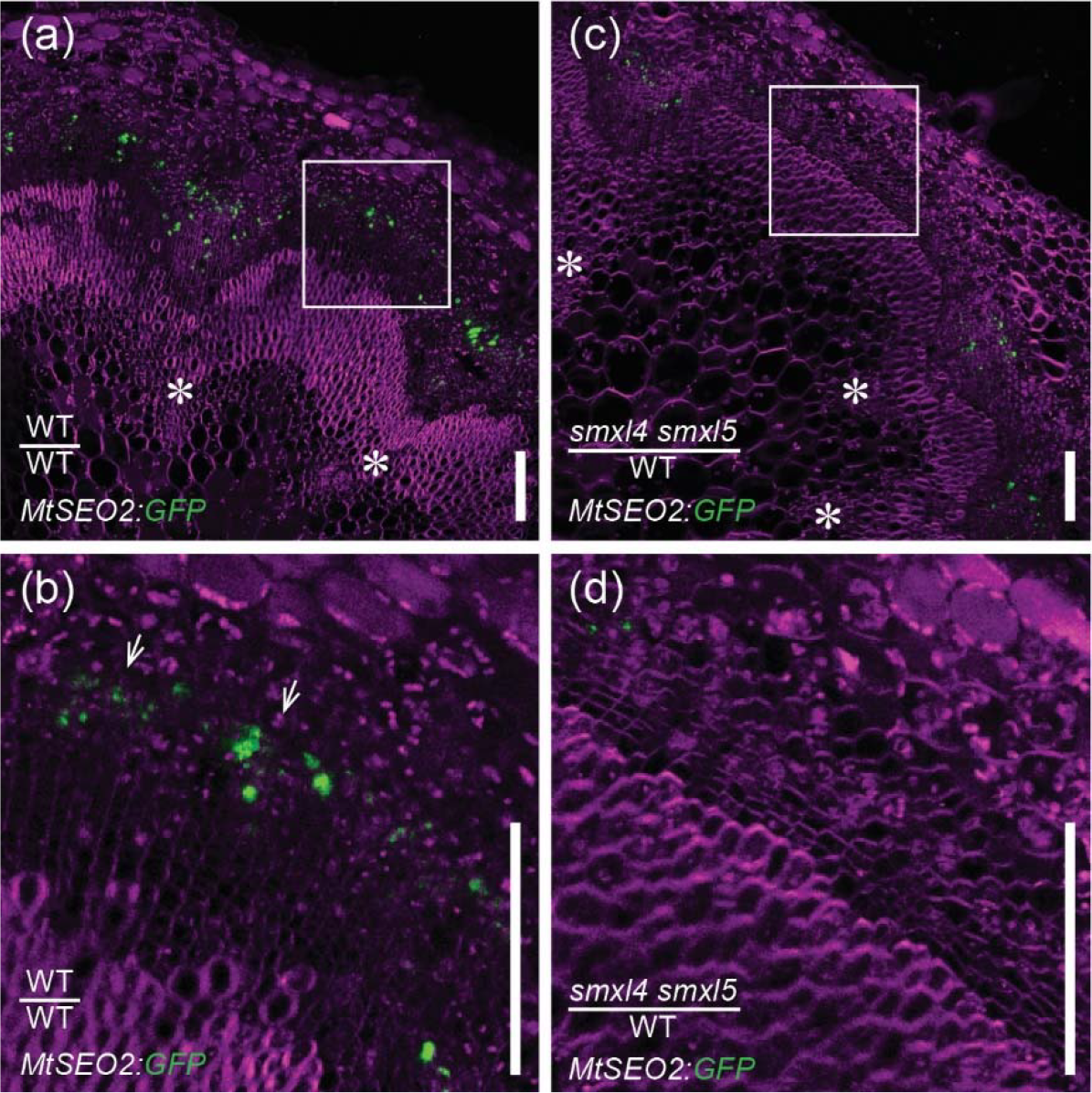
The *MtSEO2:GFP-ER* marker is not active in interfascicular regions of *smxl4;smxl5* plants (a – d) Stem base cross sections from wild type shoots grafted onto wild type roots (a, b), and from *smxl4;smxl5* shoots grafted onto wild type roots (c, d) caryying the SE-specific marker *MtSEO2:GFP-ER* were analyzed by confocal microscopy. Cell walls were counterstained by DirectRed23 (magenta). White rectangles in (a) and (c) mark interfascicular regions shown as close-ups in (b) and (d). GFP signals are shown in green. White asterisks mark interfascicular bundles. White arrows point to marker activity in secondary phloem. Scale bars represent 100 µm.

